# Inhibition of nonsense-mediated mRNA decay reduces the tumorigenicity of human fibrosarcoma cells

**DOI:** 10.1101/2023.03.28.534516

**Authors:** Sofia Nasif, Martino Colombo, Anne-Christine Uldry, Markus S. Schröder, Simone de Brot, Oliver Mühlemann

## Abstract

Nonsense-mediated mRNA decay (NMD) is a eukaryotic RNA degradation pathway that targets for degradation faulty mRNAs with premature termination codons as well as many physiological mRNAs encoding full-length proteins. Consequently, NMD functions in both, quality control and post-transcriptional regulation of gene expression, and it has been implicated in the modulation of cancer progression. To investigate the role of NMD in cancer, we knocked out SMG7 in the HT1080 human fibrosarcoma cell line. SMG7 is involved in deadenylation-coupled exonucleolytic mRNA decay, one of the two main degradation pathways in mammalian NMD. Genome-wide proteomic and transcriptomic analyses confirmed that NMD is severely compromised in these SMG7-knockout HT1080 cells. We compared the oncogenic properties between the parental, the SMG7-knockout, and a rescue cell line in which we re-introduced both isoforms of SMG7. In parallel, we tested the effect of a drug inhibiting the NMD factor SMG1 on the HT1080 cells to distinguish NMD-dependent effects from putative NMD-independent functions of SMG7. Using cell-based assays as well as a mouse xenograft tumor model, we show that the oncogenic properties of the parental HT1080 cells are severely compromised when NMD is inhibited. Molecular pathway analysis revealed a strong reduction of the matrix metalloprotease 9 (MMP9) gene expression in NMD-suppressed cells. Since MMP9 expression promotes cancer cell migration and invasion, metastasis and angiogenesis, its downregulation in NMD-suppressed cells explains, at least partially, their reduced tumorigenicity. Collectively, our findings emphasize the therapeutic potential of NMD inhibition for the treatment of certain types of cancer.

**Significance:** Nonsense-mediated mRNA decay (NMD) is a eukaryotic RNA decay pathway with reported roles in regulating cellular stress responses, differentiation, and viral defense. NMD has also emerged as a modulator of cancer progression, however, the available evidence supports both, a tumor suppressor as well as a pro-tumorigenic role for NMD. We discovered that NMD inhibition results in impaired tumorigenicity in the HT1080 human fibrosarcoma cell line and uncovered a direct correlation between NMD activity and the expression levels the pro-tumorigenic gene MMP9. Restoring MMP9 expression in NMD-suppressed cells partially improved their oncogenic properties. These results show that the tumorigenicity of the HT1080 fibrosarcoma cells relies on NMD activity and highlights the potential use of NMD inhibition as a therapeutic approach.

## Introduction

Fidelity of gene expression is essential for every living organism, which is why prokaryotic and eukaryotic cells put into play a myriad of quality control pathways to ensure that faulty RNAs and proteins are recognized and degraded. Of particular importance are RNA surveillance pathways, which not only degrade faulty RNAs but also control the levels of RNA molecules and thereby contribute to post-transcriptional gene regulation (1). In eukaryotes, the separation of transcription and translation in different cellular compartments allows for the compartmentalization and functional distinction between decay pathways aimed at degrading aberrantly processed RNAs and transcriptional byproducts, which are mainly nuclear, and decay pathways aimed at identifying and degrading RNAs with hampered functionality, most of which are cytoplasmic (2). One such cytoplasmic mRNA decay pathway is nonsense-mediated mRNA decay (NMD), which was first described as a translation-dependent mechanism responsible for the rapid decay of mRNA molecules harboring premature termination codons (PTC) (3–5). PTC-bearing mRNAs arise in eukaryotic cells as results of DNA mutations and errors occurring during transcription or splicing. In this context, NMD serves as a safeguard mechanism to prevent the production of truncated proteins that can have deleterious effects on cells. More recently, genome-wide studies revealed a much larger influence of NMD on the transcriptome of eukaryotic cells (6–11), with the latest estimates suggesting that NMD activity can modulate – directly and indirectly – the abundance of up to ~40% of the transcriptome in human cells (12). These studies showed that NMD can also target for degradation mRNAs encoding functional fulllength proteins and expanded the perception of NMD as a quality control pathway that can, additionally, exert fine post-transcriptional control of gene expression.

The mechanism of mammalian NMD has been studied in detail (reviewed in (13)). It relies on several proteins that work sequentially, and jointly, to decide the fate of the mRNA molecules. The core NMD factor is UPF1 (Up-frameshift 1), an ATP-dependent helicase that binds all mRNAs without sequence specificity (14). Selection of substrate mRNAs relies on the completion of two steps: first, activation of UPF1 by phosphorylation, mediated by the kinase SMG1 (Suppressor with morphogenetic effects on genitalia 1) and facilitated by the UPF1 interaction partners UPF2 and UPF3B, and second, recruitment of the decay effector proteins SMG5, SMG6 and SMG7 that trigger the degradation of the mRNA (13). SMG6 is an endonuclease that cleaves the mRNA in the vicinity of the stop codon (15), while SMG5 and SMG7 form a heterodimer that triggers the exonucleolytic decay of the transcript by recruiting the CCR4-NOT complex (16). There is a big overlap in the mRNAs that are targeted by the three effector proteins, suggesting that they operate with some level of redundancy (10–12, 17). In addition, it has been recently reported that the SMG5 and SMG7 proteins play a role in enabling SMG6-mediated endocleavage (12), indicating that the three NMD effector proteins work together, rather than in parallel, to trigger mRNA decay. A full understanding of the mechanistic aspects of mammalian NMD is further obstructed by its autoregulation, i.e., most of the mRNAs encoding NMD factors are substrates of the NMD pathway, and are therefore upregulated when NMD activity is compromised, attempting to restore NMD (9, 18). Additionally, it has recently been reported that NMD inhibition also results in a feedback loop, mediated by the accumulation of the NMD sensitive EIF4A2 mRNA, that boosts translation initiation. Since NMD is a translation-dependent mechanism, this feedback loop is also interpreted as trying to recover NMD activity (17). The existence of such regulatory feedback loops suggests that NMD is indispensable for cellular fitness, a notion further supported by the fact that depletion of NMD factors results in reduced viability (reviewed in (19)).

Since nonsense and frameshifting mutations account for approximately 30%of all disease-associated genetic variants, NMD’s capacity of detecting and degrading PTC-containing mRNAs makes it an important modulator of the outcome of genetic diseases (20). NMD can have opposing roles in modulating disease severity; it can be beneficial in cases where the expression of truncated proteins would have dominant negative effects, or it can worsen the disease phenotype in cases where the truncated protein would still retain some activity (20). One disease that is highly dependent on the accumulation of genetic mutations is cancer, and the link between NMD activity and cancer progression is a topic of intensive research (reviewed in (21)). There is abundant evidence supporting a pro-tumorigenic role for NMD. For instance, it has been shown that depletion of the NMD factors UPF2, UPF1 or SMG1 results in reduced *in vivo* tumor growth in a variety of cancer types, due to the strong anti-tumor immune response elicited by the NMD-deficient tumors (22–25). mRNAs containing frameshifting mutations can be translated into novel peptides, that can be recognized as foreign by the immune system and trigger an immune response. By degrading these mutant transcripts, NMD reduces the expression of neoantigens and therefore helps the tumor evade the immune attack (22–28). Further evidence that tumors benefit from high NMD activity comes from pan-cancer genomewide mutational studies that discovered that tumor suppressor genes preferentially accumulate NMD-eliciting nonsense mutations, while oncogenes are depleted of NMD-inducing mutations (29, 30). Additionally, it has been reported that the genes encoding core NMD factors are amplified in multiple types of cancers and that this is associated with higher NMD efficiency and lower cytolytic activity (31). Moreover, it was recently shown that in tumors undergoing immunotherapy, the immunosuppressive IL-6/STAT3 signaling pathway induces SMG1 levels, and NMD activity, to suppress the expression of potent frameshift-derived neoantigens (24). Another way in which NMD can promote cancer is by degrading mRNAs encoding for truncated, though still functional, tumor suppressor proteins (32–35). On the other hand, there is also evidence supporting a role for NMD as a tumor suppressor pathway. For example, it has been reported that UPF1 levels are decreased in tumor tissue, relative to normal adjacent tissue, in different cancer types (36–40). Also, a functional interplay between the tumor microenvironment and NMD has been suggested, by which cellular stressors like starvation and hypoxia reduce NMD activity and this, in turn, promotes tumorigenesis (38, 41).

To gain more insight into the molecular underpinnings of the links between NMD activity and cancer progression in a well-established tumor model, we decided to study the role of the NMD factor SMG7 in the tumorigenicity of the human fibrosarcoma cell line HT1080. We generated SMG7 knockout (KO) HT1080 cells and confirmed that they have impaired NMD. Using *in vitro* cell-based assays as well as mouse xenografts, we found that NMD-inhibited HT1080 cells display impaired tumorigenicity. Gene expression analysis revealed that NMD inhibition resulted in a greatly reduced expression of the pro-tumorigenic gene MMP9, and that the impaired tumorigenicity of the SMG7KO cells can be partially rescued by MMP9 overexpression. Our results show that in this fibrosarcoma model, NMD activity is beneficial for tumor progression and suggest that NMD inhibition would represent a promising therapeutic approach for this type of tumors.

## Results

### Lack of SMG7 results in strong NMD inhibition

The available literature showing discrepant results regarding the role of NMD in the development of tumors prompted us to investigate what is the impact of the NMD factor SMG7 in the tumorigenicity of a human fibrosarcoma model. To this end, we generated SMG7 knockout cells in the human fibrosarcoma-derived cell line HT1080 (SMG7KO) by applying CRISPR-Cas9 genome editing to introduce a gene trap (consisting of a splicing acceptor (SA), a Zeocin resistance cassette (Zeo^R^) and a strong polyadenylation signal (PA)) into the first intron of the SMG7 gene (Fig 1A) (42). In cell clones bearing the correct integration of the gene trap in both alleles, transcription of the SMG7 locus confers Zeocin resistance but no SMG7 expression, as transcription should stop at the integrated SV40 polyadenylation motif. We used western blotting (WB) to confirm the absence of both the short and the long protein isoforms of SMG7 in two independently generated SMG7KO clones (clones 11 and 12, Fig 1B) and RNA sequencing (RNAseq) to verify that there are no reads mapping to the SMG7 locus downstream of the integrated gene trap (42). To assess if SMG7KO cells have impaired NMD, we performed RNAseq and differential gene expression analysis on the SMG7KO clones 11 and 12 and on the parental cell line HT1080 (Fig 1C). The changes in the transcriptome observed in both SMG7KO clones correlate nicely with each other, suggesting that they are not specific to either clone but rather common to the loss of SMG7. In total 7258 genes were found to be differentially expressed (DEG, FDR < 0.01), with roughly half of them being up- (3522) and the other half being downregulated (3736) in SMG7KO cells. Given the role of SMG7 in RNA decay, it was surprising to find such a big fraction of downregulated genes. We reasoned that the changes in gene expression observed in SMG7KO cells are the sum of both direct and indirect (or secondary) effects of the lack of SMG7. If we restrict the analysis to the top 1000 most significant DEGs, then a clear enrichment for upregulated genes is observed, with 691 genes being up- and 301 genes being downregulated in SMG7KO cells. Among the upregulated genes, we find well validated NMD substrates like GADD45B, GAS5, RP9P and genes encoding NMD factors, as part of the NMD autoregulatory mechanism (9, 10, 43) (Fig 1C), further confirming that NMD is impaired in SMG7KO cells. In addition, we wondered if the NMD inhibition observed in SMG7KO cells could also impact their proteome. To answer this, we did a quantitative proteomic analysis on both SMG7KO clones and on parental HT1080 cells using stable isotope labeling by amino acids in cell culture (SILAC) and mass spectrometry (Fig S1A). The protein expression changes observed in both SMG7KO clones relative to HT1080 cells correlate well with each other, and among the top upregulated proteins we found many NMD factors, like UPF1, UPF2, SMG1 and SMG9 (Fig S1A). In summary, our transcriptomic and proteomic analyses document a strong NMD impairment in the SMG7KO cells.

**Figure 1.**
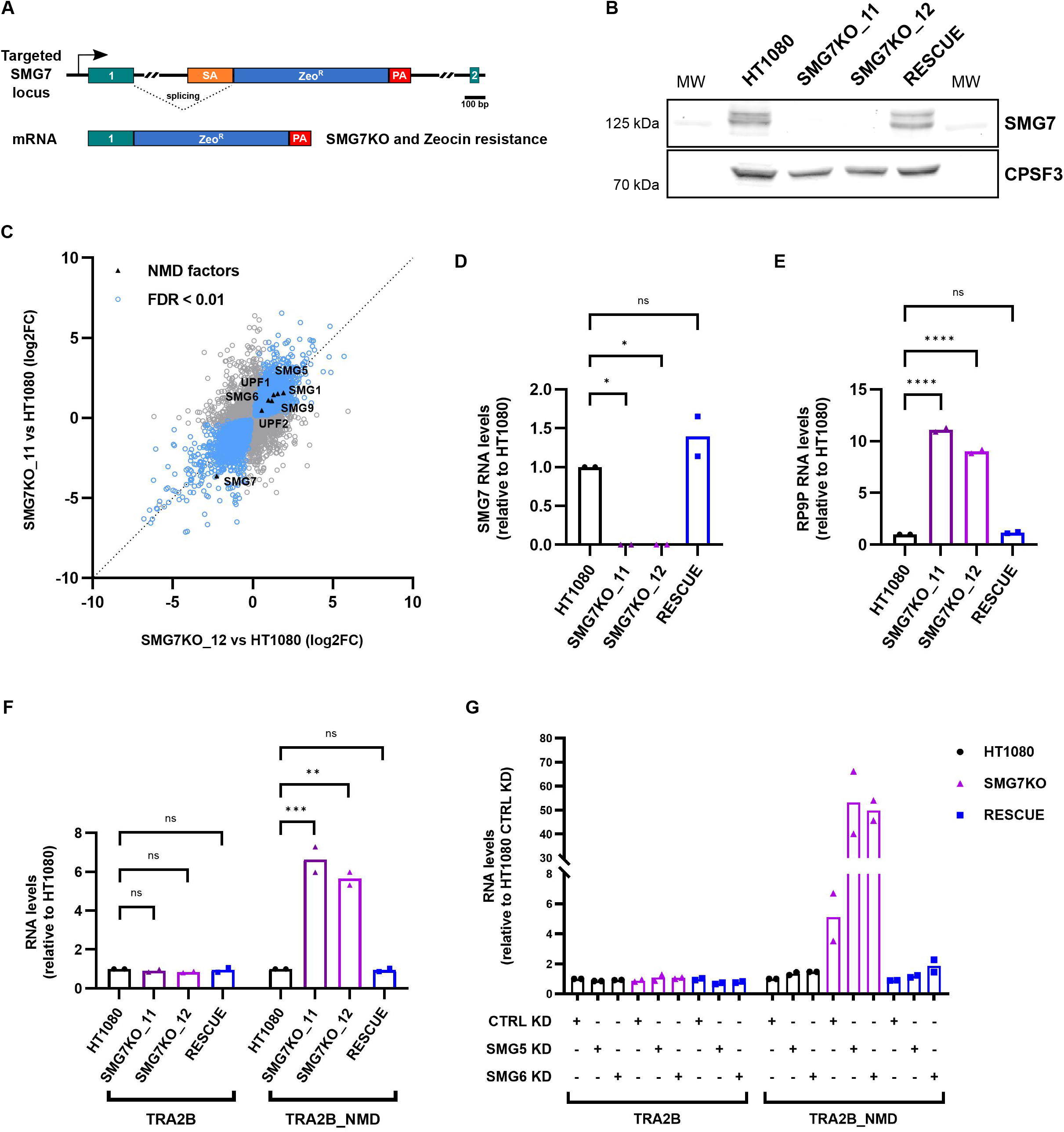
SMG7KO cells have impaired NMD. **(A)** Scheme of the genome editing approach used to generate the SMG7KO cell lines. A gene trap consisting of a splicing acceptor (SA), a Zeocin resistance cassette (Zeo^R^) and a strong polyadenylation signal (PA) was inserted into the first intron of the SMG7 gene. The first two exons of the SMG7 gene are shown in green. **(B)** Western blot showing the levels of SMG7 protein in HT1080, SMG7KO clones 11 and 12 and the RESCUE cell lines. Both isoforms of SMG7 are distinguishable, CPSF3 serves as a loading control. MW: molecular weight marker. **(C)** Scatter plot showing the correlation between the differential gene expression analyses performed on RNAseq data from SMG7KO clones 11 and 12 compared to the parental cell line HT1080. Differentially expressed genes with FDR < 0.01 are shown in blue. The genes encoding NMD factors are shown in black. **(D-F)** Relative mRNA levels, determined by RT-qPCR, normalized to ActB mRNA and 28S rRNA. The bars represent the mean values and data points represent biological replicates. Statistical significance was determined by One-way ANOVA and Dunnett’s multiple comparisons test. *ns* p > 0.05; * p ≤ 0.05; ** p ≤ 0.01; *** p ≤ 0.001; **** p ≤ 0.0001. **(G)** Relative mRNA levels, determined by RT-qPCR, in HT1080, SMG7KO_12 and the RESCUE cell lines treated with the indicated siRNAs. mRNA levels were normalized to ActB mRNA and 28S rRNA levels. Shown are the mean values, and data points represent biological replicates.

Next, we generated a RESCUE cell line in which we re-introduced both isoforms of SMG7 into the SMG7KO clone 12 (from now on termed SMG7KO, for simplicity). We used RT-qPCR to evaluate SMG7 expression and NMD activity in SMG7KO clones 11 and 12, the RESCUE cell line and in the parental cells HT1080. Analysis of SMG7 levels (Fig 1D) confirmed that SMG7KO clones do not express any detectable amounts of SMG7 mRNA and that this has been restored in the RESCUE cells, results that agree with our RNAseq data and with the SMG7 protein levels detected by WB (Fig 1B). Our RT-qPCR analysis further confirmed the specific accumulation of endogenous NMD-sensitive mRNAs in SMG7KO cells and revealed that their levels are brought back to that of the HT1080 cells in the RESCUE condition, suggesting that NMD activity has been fully restored in the RESCUE cells (Figs 1E-F and S1B-C). The model proposing that SMG5 and SMG7 proteins are required to form a heterodimer in order to trigger mRNA decay (16) was recently challenged by two publications providing evidence that SMG5 and SMG7 can have heterodimer-independent functions in NMD (12, 17). We reasoned that our cell lines provide a good framework to examine these new findings. We performed siRNA-mediated knock downs (KD) of SMG5 and SMG6 in HT1080, SMG7KO and RESCUE cells and we evaluated NMD activity by RT-qPCR. Indeed, we found that in cells lacking SMG7, depletion of SMG5 led to an even stronger NMD inhibition (Figs 1G and S1D-F), further supporting the idea that SMG5 can sustain NMD activity in the absence of SMG7. A similar picture was observed when SMG6 was depleted from SMG7KO cells, results that can be explained by the concomitant inhibition of both mRNA decay branches, as it has been reported previously (10, 12). We were reassured to see that the accumulation of NMD substrates upon depletion of SMG5 or SMG6 in the RESCUE cells was similar to that of the parental HT1080 cells, showing that the expressed recombinant SMG7 proteins are fully functional.

### NMD inhibition affects the anchorage-independent growth and migration properties of HT1080 cells

Due to their highly aggressive and invasive nature, the HT1080 cells have been used extensively as a model for studying cell invasion and migration (44–47). Given the wealth of conflicting data relating NMD and tumorigenesis (21), we decided to study if the tumorigenic potential of the HT1080 cells is affected by the loss of SMG7. First, we compared the anchorage-independent growth abilities of parental, SMG7KO and RESCUE HT1080 cells by performing soft agar colony formation assays and found a clear reduction in the number of colonies formed by SMG7KO cells, compared to both HT1080 and RESCUE cells (Fig. 2A). We did not, however, find differences in the growth rates of the cell lines under standard adherent cell culturing (Fig S2A). Similar results were obtained with another SMG7KO clone, number 11 (Fig S2B), suggesting that this is not a clone-specific trait. Additionally, we wanted to assess if the impaired ability to grow in the absence of a solid surface is a feature attributable to the lack of SMG7, or more generally to NMD inhibition. For this, we performed soft agar colony formation assays on HT1080 cells treated with a small molecule drug (NMDi) that blocks NMD by inhibiting the kinase activity of SMG1 (48, 49), and observed that the NMDi treatment greatly reduces the capacity of HT1080 cells to form colonies in soft agar (Fig 2A).

**Figure 2.**
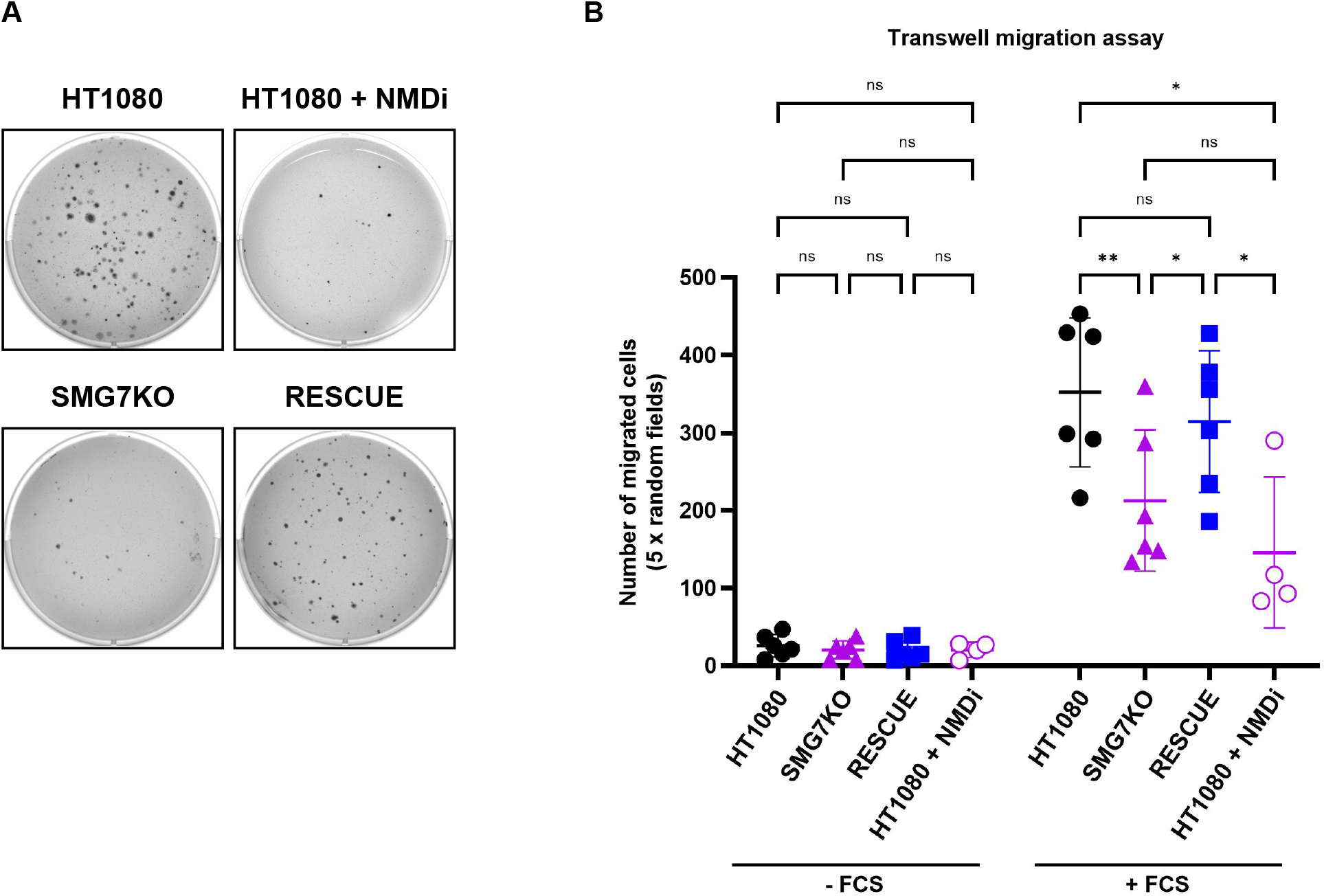
SMG7KO cells show impaired migration and anchorage-independent growth. **(A)** Soft agar colony formation assays were used to determine the anchorage-independent growth properties of HT1080, SMG7KO and RESCUE cell lines. HT1080 cells were treated with an NMD inhibitor (NMDi, 0.3 μM) as an alternative way of inhibiting NMD activity in these cells. **(B)** Scatter plot showing the number of migrated cells after transwell migration assays were performed with HT1080, SMG7KO and RESCUE cell lines. HT1080 cells were treated with 0.3 μM NMDi as an alternative way of inhibiting NMD activity in these cells. Shown are the means ± SD and data points represent biological replicates. Statistical significance was determined by Mixed-effects analysis and Tukey’s multiple comparisons test. *ns* p > 0.05; * p ≤ 0.05; ** p ≤ 0.01.

Next, we performed transwell migration assays to compare the migratory properties of the cell lines, using fetal calf serum (FCS) as a chemoattractant. We found that the number of migrating cells was significantly reduced for the SMG7KO cell line, compared to HT1080 and RESCUE cells (Fig 2B). Comparable results were obtained when HT1080 cells were treated with the NMDi, suggesting that NMD inhibition causes the impaired migratory capacity of the SMG7KO cells. In summary, our results show that the anchorage-independent growth and migratory abilities of HT1080 cells are impaired when NMD activity is compromised.

### SMG7KO cells show impaired tumorigenicity *in vivo*

The results obtained in our cell-based assays led us to hypothesize that SMG7KO cells might have difficulties in forming tumors *in vivo.* To test this, we performed mouse xenograft experiments in which parental, SMG7KO and RESCUE HT1080 cells were injected subcutaneously into immunocompromised mice and tumor growth was measured over time with a caliper. The results are summarized in figures 3A and 3B. Tumor growth was detected in all mice injected with the parental HT1080 cells, and most of these tumors reached a volume of 750 mm^3^ within about 50 days after injection (15/17, ~88% of total) (Fig 3A-B). On the contrary, tumors formed by the SMG7KO cells grew more slowly (Fig 3A). In ~35% (6/17) of the injected animals we couldn’t find evidence of tumor growth, and from the ~65% (11/17) of the cases in which tumor growth was detectable, only 4 of them (~24% of total) reached a volume of 750 mm^3^ within the timeframe of the experiment (Fig 3B). RESCUE cells were also able to form tumors *in vivo,* albeit at a slower speed than that of parental HT1080 cells (Fig 3A). There was also a fraction of RESCUE-injected mice for which tumor growth wasn’t detectable (4/17, ~24%), however, in contrast to the SMG7KO cells, a large proportion of the tumors formed by RESCUE cells reached a volume of 750 mm^3^ (10/17, ~59% of total) (Fig 3B).

**Figure 3.**
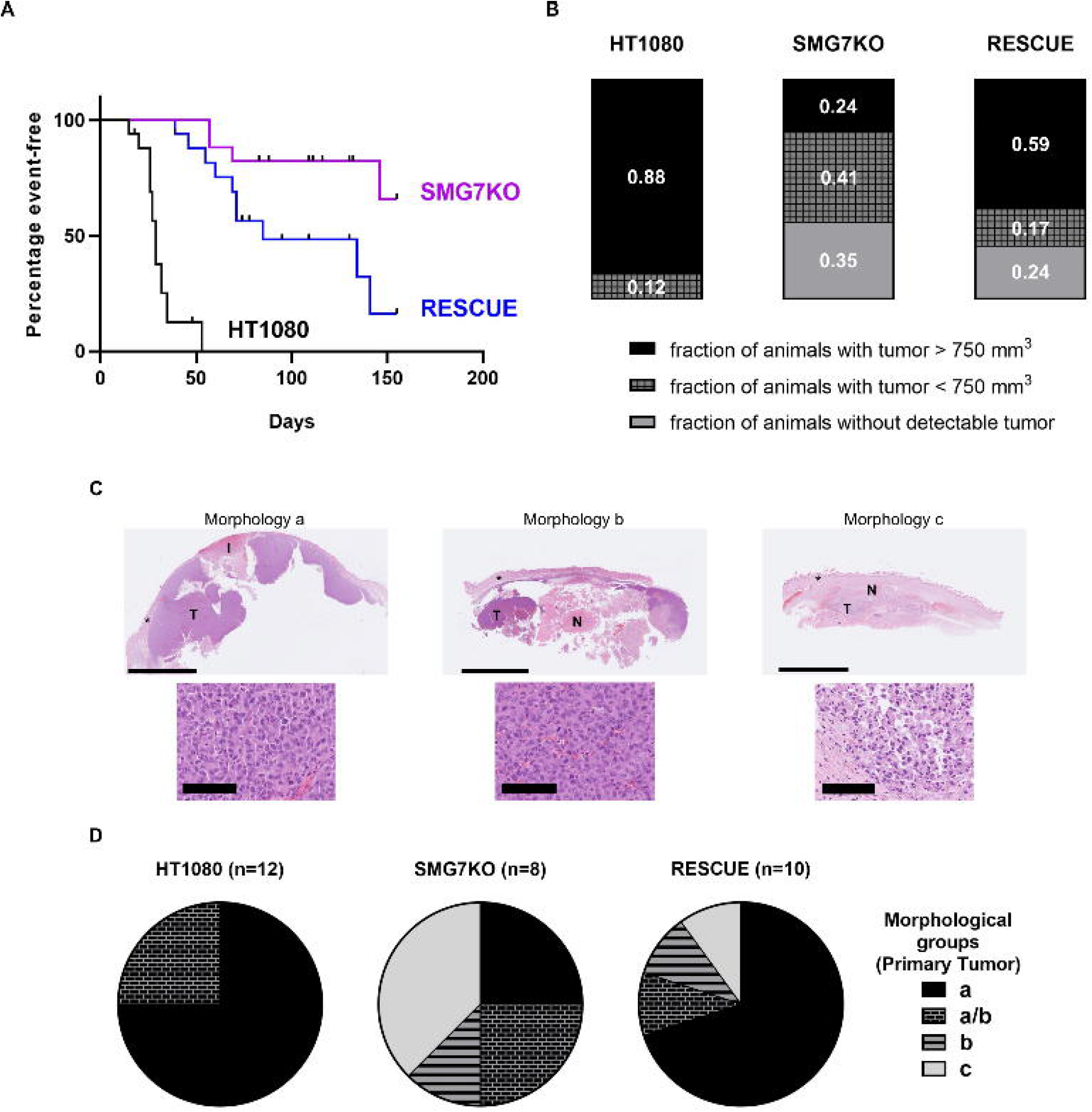
SMG7KO cells show impaired tumorigenicity *in vivo*. **(A)** Kaplan-Meier plots showing tumor progression in NSG mice with HT1080, SMG7KO and RESCUE xenografts. The endpoint (event) was defined as tumors reaching a volume of 750 mm^3^. Black ticks represent censored animals. Each group consisted of seventeen mice. Survival curves were compared using the Log-rank (Mantel-Cox) test (WT *p* < 0.0001, SMG7KO vs RESCUE *p* = 0.0149). **(B)** Stacked bars showing the variability in the take rate and tumor growth of the xenograft-injected animals during the 150 days-long outgrowth experiment. Three categories were defined: animals that did not have a detectable tumor (gray), animals with tumors smaller than 750 mm^3^ (gridded) and animals for which the tumors reached at least 750 mm^3^ (black). In total, 17 animals were injected per cell line. **(C)** Hematoxylin and eosin (H&E) staining was used for histological examination of the primary tumors. Shown are representative images of the three distinctive morphological groups termed a, b and c. The overview images (top) were taken at 0.6x and the scale bar corresponds to 5 mm. “T” denotes viable tumor tissue, “N” shows necrotic tissue, and the asterisks show the skin. “I” indicates the dermal necrosis (infarct), characteristic of tumors belonging to group a. The close-up images (bottom) were taken at 25x and the scale bar corresponds to 100 μm. The dissociation of the neoplastic cells is evident in tumors from group c. **(D)** Pie charts showing the proportion of the tumors formed by HT1080, SMG7KO and RESCUE cells that were classified into one of the three main morphological groups (a, b, or c). The fourth category labeled a/b corresponds to tumors containing features of both categories. The total number of primary tumors analyzed per cell line is indicated between parentheses.

To further characterize the tumors formed by the three cell lines, we performed a histopathological analysis of the primary tumors, which were assessed morphologically, for mitotic activity and for tumor necrosis. Based on the predominant tumor shape and the extent as well as the distribution of necrosis, the tumors demonstrated one of the following main morphologies: a) highly and densely cellular, multinodular, space-occupying mass, with frequent focally-extensive epidermal and dermal necrosis on the surface (infarct), and variable intratumoral necrosis; b) highly and densely cellular, multinodular, plaque-like more than space-occupying mass with massive central focally-extensive necrosis; c) plaque-like mass with more than 80% necrotic tumor tissue with evident dissociated neoplastic cells (Fig 3C). Some cases presented with an intermediate morphology of a) and b), characterized by a combination of histomorphological features from both groups. Morphology a) was clearly predominating in the HT1080 (75%) and RESCUE (70%) groups. For the SMG7KO group, however, none of the morphologies predominated clearly over the others, with morphology c) being the most frequently found (Fig 3D). The more plaque-like and smaller appearing tumors may correspond to slower growing neoplasia compared to the larger, multinodular, and more expansive tumors. Additionally, the extent of tumor necrosis may also potentially correspond to reduced tumor growth as the extent of necrotic lesions would be expected to increase with prolonged presence and development of the tumors. The quantification of the mitotic activity, as well as the percentage of necrosis in the primary tumors support this idea: the mitotic activity of the primary tumors was highest in the HT1080 group, lowest in the SMG7KO group and intermediate for the RESCUE group (Fig S3A). The opposite trend was observed for the quantification of tumor necrosis, which was highest in the SMG7KO group and lowest for the HT1080 group, with intermediate values for the RESCUE group (Fig S3B). Even though these differences didn’t reach statistical significance, they suggest that the tumors formed by SMG7KO cells grew more slowly than those formed by parental HT1080 cells, and that this phenotype was partially reverted in the RESCUE cells. Finally, we reconfirmed the identity of the cells present in the primary tumors by measuring SMG7 mRNA levels by qPCR (Fig S3C). Altogether, our results show that the high tumorigenicity characteristic of HT1080 cells was reduced in the SMG7KO cells, and that this attribute was partially restored in the RESCUE cells.

### NMD inhibition results in reduced MMP9 gene expression in HT1080 cells

To get more insight into the molecular mechanisms responsible for the impaired tumorigenicity of the SMG7KO cells, we performed RNAseq and differential gene expression analysis on primary tumors formed by parental, SMG7KO and RESCUE HT1080 cells (Fig 4A). The anticorrelation observed when plotting the DEGs between SMG7KO and HT1080 cells *(x* axis) vs. the DEGs between RESCUE and SMG7KO cells (*y* axis) suggests that most of the transcriptomic changes induced by the removal of SMG7 have been restored in the RESCUE cells (Fig 4A). Again, as a paradigm of NMD inhibition, we verified the accumulation of the NMD sensitive mRNAs encoding for the NMD factors UPF1, SMG1, SMG5 and SMG9 in the tumors formed by SMG7KO cells, and we also confirmed that their levels are again downregulated in the tumors established from RESCUE cells (Fig 4A).

**Figure 4.**
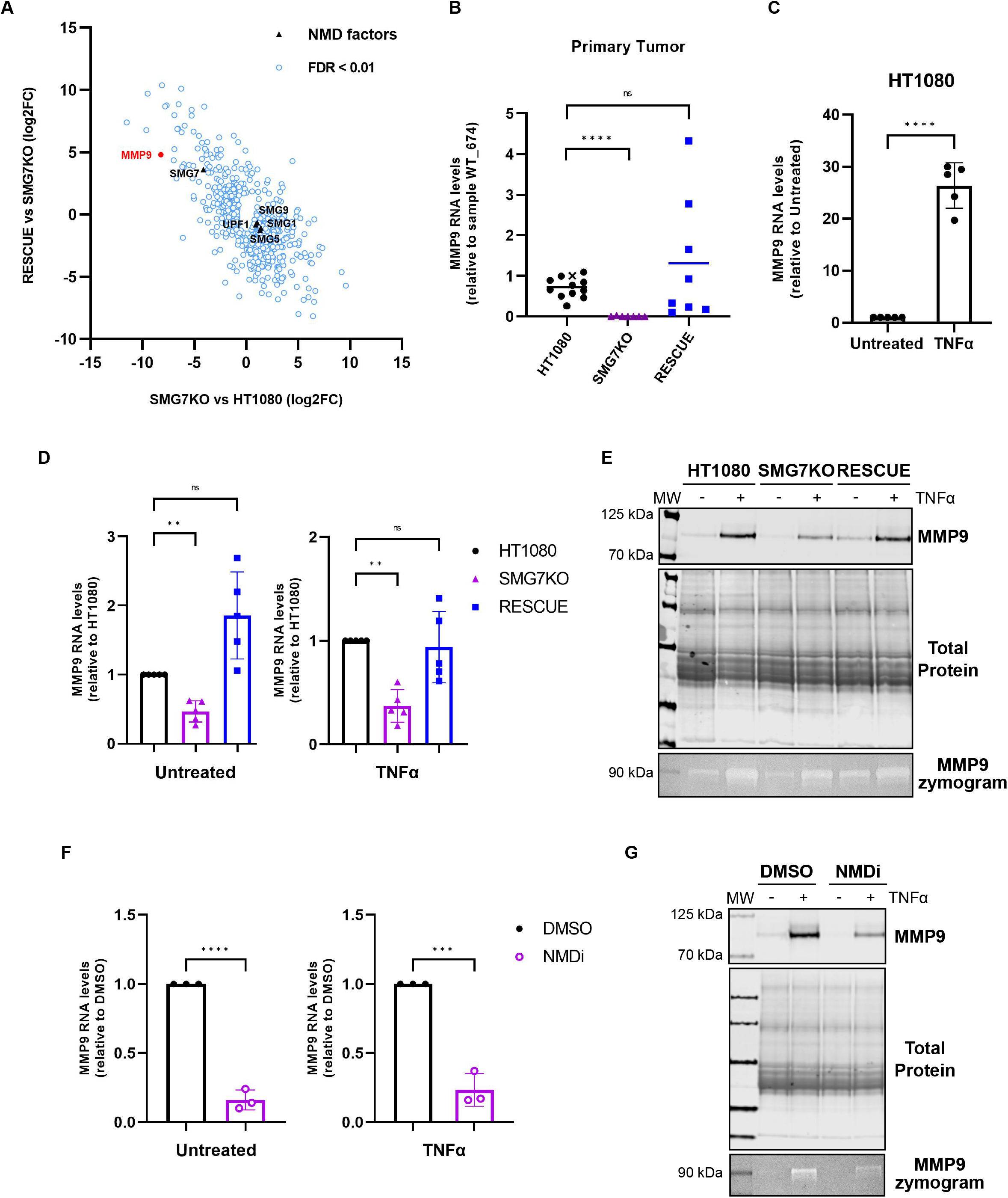
NMD inhibition leads to decreased MMP9 expression. **(A)** Scatter plot showing the anti-correlation between the differential gene expression analyses performed on RNAseq data from primary tumors formed by HT1080 vs SMG7KO cells (*x* axis) and those formed by RESCUE vs SMG7KO cells (*y* axis). Only differentially expressed genes with an FDR <0.01 are plotted (in blue). The genes encoding NMD factors are shown in black and the MMP9 gene is shown in red. **(B)** MMP9 gene expression in primary tumors. Relative MMP9 mRNA levels were determined by RT-qPCR and normalized to ActB mRNA. Shown are the means and each data point represents a different primary tumor, the values are expressed relative to the tumor sample HT1080_674, marked with a “x”. Statistical significance was determined by unpaired two-tailed t-test. *ns* p > 0.05; **** p ≤ 0.0001. **(C)** Relative MMP9 mRNA levels in HT1080 cells untreated or treated with TNFα (25 ng/ml). MMP9 expression was determined by RT-qPCR, normalized to ActB mRNA and 28S rRNA. Shown are the mean values ± SD and data points represent biological replicates. Statistical significance was determined by unpaired two-tailed t-test. **** p ≤ 0.0001. **(D)** Relative MMP9 mRNA levels in HT1080, SMG7KO and RESCUE cells untreated or treated with TNFα (25 ng/ml). MMP9 expression was determined by RT-qPCR, normalized to ActB mRNA and 28S rRNA. Shown are the means ± SD and data points represent biological replicates. Statistical significance was determined by One-way ANOVA and Dunnett’s multiple comparisons test. *ns* p > 0.05; ** p ≤ 0.01. **(E)** Analysis of MMP9 protein expression and activity. Western blot was used to analyze the levels of MMP9 protein present in conditioned medium collected from HT1080, SMG7KO and RESCUE cells untreated or treated with TNFα (25 ng/ml). Total protein staining serves as a loading control. Lower panel: gelatin zymography was used to detect the gelatinolytic activity of secreted MMP9. **(F)** Relative MMP9 mRNA levels in HT1080 cells treated with DMSO or with the NMD inhibitor (NMDi, 0.3 μM) in the absence or presence of TNFα (25 ng/ml). MMP9 expression was determined by RT-qPCR, normalized to ActB mRNA and 28S rRNA. Shown are the mean values ± SD and data points represent biological replicates. Statistical significance was determined by unpaired two-tailed t-test. *** p ≤ 0.001; **** p ≤ 0.0001. **(G)** Analysis of MMP9 protein expression and activity. Western blot was used to analyze the levels of MMP9 protein present in conditioned medium collected from HT1080 cells treated with DMSO or with 0.3 μM NMDi in the absence or presence of TNFα (25 ng/ml). Total protein staining serves as a loading control. Lower panel: gelatin zymography was used to detect the gelatinolytic activity of secreted MMP9.

We then used the Ingenuity Pathway Analysis (IPA) software to perform a functional analysis on the RNAseq data of the primary tumors. Using the “Downstream effects” analysis tool, which uses the gene expression changes in the dataset to predict which biological processes or functions might be activated or inhibited, we found categories like “Invasion of cells”, “Advanced malignant tumor”, “Invasive tumor”, and “Metastasis” to be inhibited in the tumors formed by SMG7KO cells (z-scores < −2) (Fig S4A), a result that is consistent with the observed phenotype. In addition, we implemented a “Regulator effects” analysis to get insight into the upstream regulators and the DEGs that might be driving the predicted downstream effects. This analysis resulted in only one causal network with a positive consistency score (a measure of how causally consistent and densely connected the network is) that connects 13 DEGs that are predicted to contribute to the inhibition of the categories “Migration of cells”, “Invasion of cells” and “Metastasis” (Fig S4B). We also used the “Comparison” tool from the IPA software to study which DEGs and downstream affected diseases or functions are shared between the RNAseq data from the primary tumors and that of the cells in culture. We found that the gene expression changes observed in the cultured cells also suggested the inhibition of categories like “Invasion of cells”, “Metastasis” and “Advanced malignant tumor” in the absence of SMG7 (Fig S4C).

Among the DEGs reported to affect these downstream categories, we found Matrix Metallopeptidase 9 (MMP9; also known as 92 kDa Gelatinase) to be a very interesting candidate for further study. MMP9 9 is a proteinase that degrades type IV and V collagens and its activity has been suggested to play a role in extracellular matrix remodeling during tumorigenesis (50–52). MMP9 is among the most downregulated genes in the tumors formed by SMG7KO cells and its expression is reestablished in the RESCUE tumors (Fig 4A), results that were further validated by analyzing MMP9 mRNA levels in the primary tumors by RT-qPCR (Fig 4B). The transcriptomic analysis from the cells in culture also showed that MMP9 gene expression is downregulated in the SMG7KO cells (SMG7KO_12 log2FC: −3.42; SMG7KO_11 log2FC: −2.29). The expression level of MMP9 in the cultured cells is, however, quite low, and we therefore used TNFα stimulation, which induces MMP9 expression by about 25-fold (Fig 4C). We compared the levels of MMP9 mRNA in HT1080, SMG7KO and RESCUE cells by RT-qPCR and found that, either untreated or after TNFα stimulation, SMG7KO cells consistently expressed less MMP9 mRNA than HT1080 and RESCUE cells (Fig 4D). We confirmed that the downregulation in MMP9 mRNA levels also results in less secreted MMP9 protein and, consequently, less gelatinolytic activity detected in conditioned media from SMG7KO cells (Fig 4E). To understand if the observed downregulation of MMP9 levels can be attributed to NMD inhibition, we analyzed the expression levels and activity of MMP9 in HT1080 cells treated with the NMDi and found that this alternative way of inhibiting NMD also resulted in downregulation of MMP9 expression (Figs 4F-G and S4D). Furthermore, we also observed diminished MMP9 expression in HT1080 cells depleted of the core NMD factor UPF1 (Fig S4E-F). Finally, we were wondering if the reduced expression of MMP9 under NMD inhibition stems from a failure of these cells to respond to TNFα. To answer this question, we analyzed the expression of the two TNFα early-response genes A20 and NFKBIA (53) after 1 hour of TNFα stimulation. We didn’t find differences in the induction of these genes in UPF1-depleted cells, compared to a control knockdown in HT1080 cells (Fig S4G), or between HT1080, SMG7KO, and RESCUE cells (Fig S4H). We therefore conclude that the failure to upregulate MMP9 cannot be explained by a general insensitivity to TNFα in NMD-inhibited cells. In summary, our results show that NMD inhibition results in reduced expression of the MMP9 gene in HT1080 cells.

### MMP9 overexpression improves the anchorage-independent growth and migration capacities of SMG7KO cells

MMP9 expression has been suggested to directly impact cell invasion, migration, and metastasis in a variety of cell lines and model organisms (51, 54–57). Given the direct implication of MMP9 in these essential processes for tumorigenicity, we wanted to test if increasing MMP9 expression in SMG7KO cells could rescue some of the phenotypes we described above. We genome-edited the SMG7KO cells to overexpress a C-terminally FLAG-tagged version of MMP9, a cell line we refer to as MMP9OE. We confirmed that the recombinant protein is expressed in high quantities, it is secreted into the conditioned medium and it is active in degrading gelatin (Fig 5A). We next compared the migration properties of SMG7KO and MMP9OE cells by doing transwell migration assays and found that MMP9OE cells show increased migration compared to SMG7KO cells (Fig 5B). We also performed colony formation assays and found that MMP9 overexpression partially rescues the soft-agar growth impairment of the SMG7KO cells (Fig 5C). Overall, we found that the defects in anchorage-independent growth and migration capacities caused by NMD inhibition in the HT1080 cells can be partially rescued by overexpression of MMP9.

**Figure 5.**
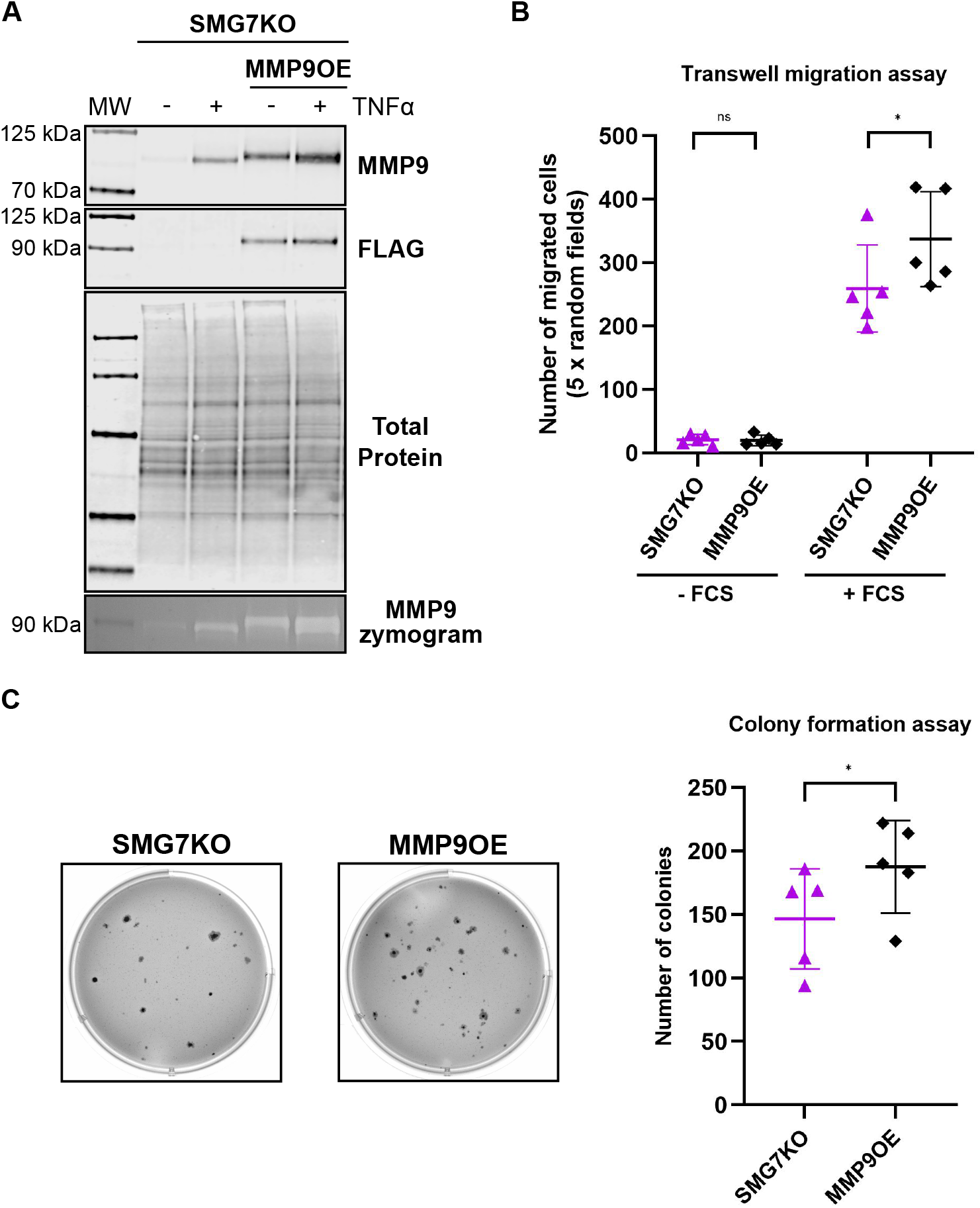
MMP9 overexpression improves the anchorage-independent growth and migratory properties of SMG7KO cells. **(A)** Analysis of MMP9 protein expression and activity. Western blot was used to analyze the levels of MMP9 protein present in conditioned medium collected from SMG7KO and MMP9OE cells untreated or treated with TNFα (25 ng/ml). Anti-FLAG antibody detects only the recombinant MMP9. Total protein staining serves as a loading control. Lower panel: gelatin zymography was used to detect the gelatinolytic activity of secreted MMP9. **(B)** Scatter plot showing the number of migrated cells after transwell migration assays were performed with SMG7KO and MMP9OE cell lines. Shown are the mean values ± SD and data points represent biological replicates. Statistical significance was determined by paired two-tailed t-tests. *ns* p > 0.05; * p ≤ 0.05. **(C)** Soft agar colony formation assays were used to compare the anchorage-independent growth properties of SMG7KO and MMP9OE cell lines. The total number of colonies formed by each cell line was quantified and the scatter plot on the right shows the mean values ± SD and data points represent biological replicates. Statistical significance was determined by paired two-tailed t-test. * p ≤ 0.05.

## Discussion

In mammalian cells, NMD is an essential RNA degradation pathway, with a continuously growing list of reported roles in cellular physiology (19). Among these, evidence for different impacts of NMD on tumor development has emerged over the past years, with examples ranging from NMD benefitting tumor growth to NMD inactivation being specifically associated with tumor tissue (21). Here, we used the human fibrosarcoma cell line HT1080 to investigate the roles of the NMD factor SMG7 in tumorigenesis. Using CRISPR-Cas9-based approaches, we generated SMG7KO cells and its RESCUE counterpart, and we confirmed that NMD activity is strongly inhibited in the former and restored in the latter (Figs. 1 and S1). We found that the NMD-inhibited SMG7KO cells show impaired cell migration and clonogenic abilities (Figs. 2 and S2), and that they display severe delays when challenged to form tumors *in vivo* (Figs. 3 and S3). Gene expression analysis showed that NMD-inhibited cells fail to upregulate the expression of the pro-tumorigenic gene MMP9 both *in vivo* and in cell culture (Figs 4 and S4). Finally, we showed that restoring MMP9 expression in SMG7KO cells partially rescues their clonogenicity and migration properties (Fig 5). Collectively, our work shows in this fibrosarcoma model, that the tumor cells depend on NMD to exert their tumorigenic properties.

Since the realization that NMD targets for degradation not only faulty mRNAs but also mRNAs with full coding potential (for a comprehensive list see (58, 59)), there has been a longstanding quest to identify cellular functions and pathways that are susceptible to changes in NMD activity (19, 60). Efforts to answer these questions in mammalian cells included RNAi-mediated transient KD experiments (10, 61, 62) and, more recently, generation of complete KOs of NMD factors (12, 17, 63, 64). The commonly used strategy for generating KOs by introducing Cas9-driven indels in the coding region of a gene, which will in most cases lead to a frameshift and a premature termination codon, relies on the NMD pathway to destroy the mutant mRNA. This editing strategy therefore might not be well suited when the target gene encodes an NMD factor, because NMD inhibition would result in stabilization of the mutant mRNA, allowing for translation of truncated versions of the protein that could have deleterious or unforeseen effects on cells. By contrast, the CRISPR-Trap approach (42) overcomes this caveat and proved to be very efficient in allowing the stable and complete removal of the SMG7 mRNA and protein. This KO of SMG7, in turn, uncovered the profound effect that SMG7 depletion has on the transcriptome of human cells, which contrasts with the mild phenotypes obtained previously for SMG7 KD experiments (9, 10, 12). This discrepancy can have two non-mutually exclusive explanations: on the one hand, the temporal distinction between transient KDs and stable KOs may account for the differences in the magnitude and the number of deregulated genes; on the other hand, it is conceivable that low residual levels of SMG7 can still sustain NMD activity in transient KD experiments and that this can only be surpassed when SMG7 is completely ablated from the cells. Our work reinforces the recent findings that SMG5 and SMG7 can have heterodimer-independent roles in NMD (12, 17). Specifically, we showed that in the absence of SMG7, SMG5 is still engaged in sustaining NMD activity, as evidenced by the strong upregulation of NMD substrates when SMG5 was depleted from SMG7KO cells. The extent to which NMD substrates accumulated in SMG7KO/SMG5KD and SMG7KO/SMG6KD conditions was remarkably similar, suggesting that both combinations lead to a comparable level of NMD inhibition. These results agree with the notion that the concomitant depletion of SMG5 and SMG7 also affects the activity of the endonucleolytic SMG6-mediated mRNA decay, as recently reported (12), and with the fact that the level of NMD inhibition achieved by the co-depletion of SMG5 and SMG7 is not further increased by the depletion of all three NMD effectors simultaneously (12, 17).

It is well established that the accumulation of somatic genetic alterations drives the development of cancer (65). Consequently, cancer cells express more PTC-bearing mRNAs compared to non-cancerous cells because of such frameshifting or nonsense mutations. The impact of these mutations on cancer fitness depends not only on whether they are driver mutations occurring in oncogenes or tumor suppressor genes, or passenger mutations that do not confer a growth advantage to the tumor cells, but also on whether they are targeted by NMD and, if they are, to which extent. Tumor evolution is a Darwinian process, and genome-wide studies have highlighted how the selective pressure has shaped the mutational landscape and NMD activities of tumor cells to benefit cancer development. Pancancer studies have shown that tumor suppressor genes preferentially accumulate NMD-eliciting mutations, whereas oncogenes are under purifying selection against mutations that trigger NMD (29, 30). Furthermore, it has been reported that the core NMD factor genes are amplified in diverse tumor types, and that this correlates with increased NMD efficiency and lower cytolytic activity (31), suggesting that enhanced NMD activity might be beneficial for tumor growth. In addition, NMD activity can be instrumental for cancers to evade the immune response. mRNAs containing frameshift mutations, if translated, can give rise to novel peptides that can be recognized as neoantigens by the immune system and trigger an anti-tumor response. However, NMD can limit the expression of neoantigens by degrading the mutant mRNAs (24, 26, 66). It has been reported that cancers with a high frequency of NMD-insensitive frameshift mutations show increased immune infiltration and are associated with a better outcome after treatment with immune checkpoint inhibitors (27, 28). Consistent with this, NMD inhibition results in strong immune responses and reduced tumor growth *in vivo* (22–25). Our findings that NMD inhibition results in decreased tumorigenicity in the HT1080 fibrosarcoma model is fully in line with these reports. We experimentally validated the *in vivo* tumorigenic impairment for HT1080 cells lacking SMG7, and our cell-based *in vitro* assays showed that the reduced migration and clonogenic capacities characteristic of the SMG7KO cells could be reproduced in the parental cells by inhibiting the SMG1 kinase, strongly suggesting that the impaired tumorigenicity might be attributed to NMD inhibition rather than to NMD-independent functions of the SMG7 protein (67). The necessity to perform our xenograft experiments in NOD scid gamma (NSG) mice, which is one of the most immunocompromised mouse strains, precludes us from assessing if the implanted SMG7KO cells can trigger a stronger immune response than that of parental or RESCUE cells. However, the IPA analysis conducted on the RNAseq dataset of the tumors revealed that the tumors formed by SMG7KO cells showed changes in gene expression that are consistent with increased inflammatory processes, results that agree with previous findings (68, 69).

Our work uncovered a direct correlation between NMD activity and MMP9 expression levels, both *in vivo* and *in vitro,* and MMP9 expression has been associated with the malignant phenotype of many different cancer types (50, 51, 54–56, 70–72). Consistently, we could show that the migration and anchorage-independent growth deficits in SMG7KO cells could be partially rescued by overexpressing recombinant MMP9. Considering the reported roles for MMP9 in the spreading of tumor cells, we hypothesize that NMD inhibited cells could show impaired metastatic capacity, and the use of experimental metastasis models (73) will be fundamental to understand if NMD inhibition could be used to limit the spreading of cancer cells to other organs. It remains to be investigated what is the molecular mechanism by which NMD inhibition results in MMP9 downregulation. MMP9 expression has been shown to be modulated by epigenetic, transcriptional, and post-transcriptional mechanisms (74–79). Therefore, further experiments are needed to completely elucidate this missing molecular link. In addition, the forced overexpression of MMP9 in SMG7KO cells resulted only in a partial rescue of the anchorage-independent growth phenotype. We therefore envision that other de-regulated genes might also contribute to the observed phenotype. Our IPA analysis on tumor samples revealed many genes that could potentially impact the tumorigenicity of HT1080 cells, including ICAM1 (80, 81), ANGPTL2 (82) or LTF (83), and follow up studies are required to explore the potential contributions of these de-regulated genes.

Collectively, our results provide insights into the molecular mechanisms by which NMD activity benefits tumorigenicity of a classical cancer model system. Together with recent reports providing evidence for NMD’s beneficial role in the development of a variety of different cancer types (22–25), our findings suggest that in many cases, inhibition of NMD might represent a promising strategy to reduce tumor growth and prevent metastasis.

## Materials and Methods

### Cell lines and cell culture

HT1080 cells (CCL-121™, ATCC®), and all cell lines derived from them, were cultured in Dulbecco’s Modified Eagle Medium (DMEM) supplemented with 10% fetal calf serum (FCS), 100 U/ml penicillin and 100 μg/ml streptomycin (P/S) at 37 °C under 5% CO2. TNFα treatment (25 ng/ml rh TNF-alpha, STEMCELL Technologies) was performed in DMEM without FCS for 20 hours, unless otherwise stated. NMD inhibition (NMDi) was achieved by supplementing the medium with 0.3 μM of a SMG1 inhibitor (hSMG1-inhibitor 11e, PC-35788 ProbeChem®). When NMDi and TNFα treatments were combined, cells were first treated for 24 hours with NMDi and then TNFα was added for 20 hours before harvesting.

SMG7KO cells were generated by introducing a gene trap into the first intron of the SMG7 gene, via CRISPR-Cas9 genome editing, as described in (42). Single cell-derived clones were picked after Zeocin selection and two of them, SMG7KO_11 and SMG7KO_12, were used here.

The RESCUE cell line was generated by re-introducing both isoforms of SMG7 in the SMG7KO_12 clone, using the AAVS1 Safe Harbor Targeting System (GE622A-1-SBI, System Biosciences). In the first round of genome editing, a construct encoding the N-terminally 3xFlag-tagged short isoform of SMG7 (3xFlag-SMG7short) and a puromycin resistance gene was used. After transfection, cells were selected for 10 days with puromycin (2 μg/ml) and single cell-derived clones were picked. The clones were analyzed by screening for SMG7 mRNA expression using qPCR, and expression of 3xFlag-SMG7short was confirmed by western blotting (WB). Next, a clone expressing 3xFlag-SMG7short was subjected to a new round of genome editing, in which we introduced a N-terminally HA-tagged SMG7long expression cassette along with a blasticidin resistance gene. After selection for 10 days with 6 μg/ml blasticidin, single cell-derived clones were picked and analyzed by WB to confirm the expression of both SMG7 isoforms.

To generate the MMP9 overexpressing (MMP9OE) cell line, the MMP9 coding sequence was amplified from cDNA of one of the tumors formed by HT1080 cells (Tumor WT_674) and subcloned into a pcDNA5-derived expression vector, in which the hygromycin resistance gene was replaced by a PGK promoter followed by a puromycin selection cassette. Subcloning into this vector also introduced a 3xFlag tag in the C-terminus of MMP9. This plasmid was then used to generate a cell line that constitutively expresses MMP9-3xFlag. For this, SMG7KO_12 cells were transfected with the expression plasmid and selected for 10 days with 1 μg/ml puromycin. Stably transfected cells were kept as a pool, and MMP9 expression and gelatinolytic activity was confirmed by performing WB with anti-MMP9 and anti-Flag antibodies and gelatin zymography, respectively.

### Cellular transfection

Plasmid transfections to generate stable cell lines were performed with TransIT®-LT1 transfection reagent (Mirus). 2×10^5^ cells per well were seeded in 6-well plates the day before transfection, transfection complexes containing 3 μl of the transfection reagent per μg of transfected DNA were prepared in Opti-MEM™ (Gibco) medium, incubated 20 minutes at room temperature and then added onto the cells. The medium was exchanged on the following day and antibiotic selection started on the third day after transfection.

For siRNA transfections, 2×10^5^ cells per well were seeded into a 6-well plate and transfections were performed using the Lullaby transfection reagent (OZ Biosciences). Transfection complexes were prepared in Opti-MEM™ (Gibco) medium and consisted of 5.6 μl of the transfection reagent and 22 nM siRNA in a final volume 2 ml/well. Cells were transfected twice with the siRNAs, on the first and third days after seeding, and harvested one day after the second transfection.

### Protein analysis by western blotting

Cellular protein levels were analyzed by lysing cells in RIPA buffer (10 μl per 1×10^5^ cells; 10 mM Tris-HCl pH 8.0, 1 mM EDTA, 1% Triton X-100, 0.5% sodium deoxycholate, 0.1% SDS, 150 mM NaCl, supplemented with protease and phosphatase inhibitors), followed by WB. Typically, lysate corresponding to 2×10^5^ cell equivalents was mixed with loading buffer (NuPAGE™ LDS Sample Buffer, Thermo Fisher Scientific) containing 50 mM DTT, heated for 10 minutes at 70 °C, and loaded onto NuPAGE™ 4-12% Bis-Tris gels (Thermo Fisher Scientific). Transfer to nitrocellulose membranes (iBlot™2 Transfer Stacks, Thermo Fisher Scientific) was done with the iBlot2 Dry Blotting System (Thermo Fisher Scientific), using the P0 standard program. After blocking for 1 hour at room temperature with 5% BSA (in 20 mM Tris, 150 mM NaCl, 0.1% (v/v) Tween 20, pH 7.4), membranes were probed with the specified primary antibodies (diluted in blocking solution), either for 2 hours at room temperature or overnight at 4 °C. We used fluorophore-coupled secondary antibodies and fluorescent signals were detected using the Odyssey Infrared Imaging System (LI-COR Biosciences). The antibodies used in this study are listed in table S1.

For studying the levels of secreted proteins, 2 ml of conditioned media were collected from nearly confluent 6-well plates, centrifuged for 10 minutes at 16’000 g and the cleared supernatant was transferred to a new tube. Half of the collected material was stored at −80 °C, while the other half was subjected to protein precipitation with acetone. Precipitated proteins were resuspended directly in loading buffer (NuPAGE™ LDS Sample Buffer, Thermo Fisher Scientific) and loaded onto NuPAGE™ 4-12%Bis-Tris gels (Thermo Fisher Scientific). After proteins were transferred to nitrocellulose membranes, they were stained with Revert700 total protein stain (LI-COR Biosciences) and imaged using the Odyssey Infrared Imaging System (LI-COR Biosciences), for normalization. After the total protein stain was removed, the WB protocol was followed as described above.

### RNA extraction and mRNA expression analysis by RT-qPCR

Total RNA from cultured cells was isolated using the guanidium thiocyanate-phenol-chloroform extraction protocol followed by isopropanol precipitation as described in (84). Residual DNA was removed with the TURBO DNA-free™ kit (Thermo Fisher Scientific) and RNA concentration was measured by NanoDrop 2000 spectrophotometer (Thermo Fisher Scientific). Next, 1 μg of RNA was reverse transcribed using random hexamers and the AffinityScript Multi-Temp reverse transcriptase (Agilent) following the manufacturer’s instructions. cDNA levels were measured by quantitative realtime PCR using Brilliant III Ultra-Fast SYBR Green qPCR mix (Agilent) and 24 ng of cDNA, in 15 μl reactions. Cycling and fluorescence acquisition were done with the Rotor-Gene Q (Qiagen). RT minus reactions were run in every experiment to control for DNA contamination. All the assays used in this study have an amplification efficiency of 1, allowing the use of the ΔΔCt method for relative quantitation. The primer sequences used for qPCR can be found in table S1.

To extract total RNA from the tumor samples, the tissue was first lysed and homogenized by dual centrifugation at 2500 rpm, −5 °C for 4 minutes using the ZentriMix 380R (Hettich) and RNA was purified using the GeneElute™ Mammalian Total RNA Midiprep Kit (Sigma Aldrich) following the manufacturer’s instructions, including the in-column DNAse digestion step. The extracted RNA was then reverse transcribed and cDNAs were quantified by qPCR as described above. For measuring SMG7 expression levels in the tumor samples, a SMG7 TaqMan qPCR assay was used, because it is specific for detecting SMG7 of human origin. For this, the Brilliant III Ultra-Fast qPCR mix (Agilent) was used.

### RNAseq analysis

The RNAseq analysis of the HT1080, SMG7KO_11 and SMG7KO_12 cell lines has been described in detail in (42).

For the transcriptomic analysis of the tumor samples, libraries were prepared with the Illumina TruSeq Stranded Total RNA library kit, including rRNA depletion. The NovaSeq6000 instrument was used to produce 50 bp paired-end reads, obtaining on average 25 million reads per sample. Two independently grown tumors were analyzed per injected cell line. The reads were mapped to the human (hg38) and mouse (mm10) genomes, using STAR. On average, ~80% of the reads mapped to the human genome. Counting of the reads and differential gene expression analysis was done using featureCounts and DESeq2, respectively. Gene annotations were taken from biomaRt.

### Proteomic analysis by SILAC and MS/MS

HT1080, SMG7KO_11 and SMG7KO_12 cells were grown for 8 cell doublings in SILAC labelling medium (85). Each cell line was labelled once with light- and once with heavy-amino acids. After harvesting, cells were lysed in lysis buffer (8 M urea in 100 mM Tris-HCl pH 8.0) supplemented with protease inhibitors. Next, the light- and heavy-labelled protein lysates of the cell lines to be compared were mixed in a 1:1 ratio, reduced, alkylated and precipitated with cold acetone. Finally, the purified protein mixtures were resuspended in loading buffer (NuPAGE™ LDS Sample Buffer, Thermo Fisher Scientific) and 40 μg of protein mix per lane were loaded on a NuPAGE™ 4-12% Bis-Tris gel (Thermo Fisher Scientific). After electrophoresis, the gel was stained with Coomassie Brilliant Blue and the gel lanes were cut into 22 slices. The gel pieces were reduced, alkylated and digested with trypsin (86). The digests were analyzed by liquid chromatography (LC)-MS/MS (PROXEON coupled to a QExactive HF mass spectrometer, ThermoFisher Scientific) with one injection of 5 μl digests. Peptides were trapped on a μPrecolumn C18 PepMap100 (5μm, 100 Å, 300 μm × 5mm, ThermoFisher Scientific, Reinach, Switzerland) and separated by backflush on a C18 column (5 μm, 100 Å, 75 μm × 15 cm, C18) by applying a 40-minute gradient of 5% acetonitrile to 40% in water, 0.1% formic acid, at a flow rate of 350 nl/min. The Full Scan method was set with resolution at 60,000 with an automatic gain control (AGC) target of 1E06 and maximum ion injection time of 50 ms. The data-dependent method for precursor ion fragmentation was applied with the following settings: resolution 15,000, AGC of 1E05, maximum ion time of 110 milliseconds, mass window 1.6 m/z, collision energy 27, under fill ratio 1%, charge exclusion of unassigned and 1+ ions, and peptide match preferred, respectively.

The mass spectrometry data was searched with MaxQuant (87) software version 1.5.4.1 against the Human Swiss-Prot (88) database (release 2016_04). Heavy labels were given as arg (R10) and lys (K8) with a maximum of three labeled amino acid in a peptide; digestion mode was set to Trypsin/P, allowing for a maximum of 2 missed cleavages; first search peptide tolerance was set to 10 ppm, and MS/MS match tolerance to 20 ppm; allowed variable modifications were Oxidation (M) and Acetyl (Protein N-term), with a maximum of 3 modifications allowed per peptide; Carbamidomethyl (C) was set as a fixed modification. Match between runs was enabled between identical gel bands and their direct neighbors with default parameters. Each pair of ratio and reversed ratio samples were analyzed to identify protein groups behaving inconsistently. This was done by plotting the log2 ratio versus the log2 inverse ratio and identification of regression outliers from absolute Studentized residuals (R MASS package (89)) > 5.209 (0.9999999th quantile of the corresponding Student distribution). Protein groups flagged as outliers were not further considered. The strength of the pair correlation was then estimated using the R duplicateCorrelation package (90), and as suggested, used as parameter in the empirical Bayes approach (moderated t-statistics (91)) used to evaluate differential expression between the two groups. Resulting *p*-values were adjusted for multiple testing by the Benjamini and Hochberg false discovery rate-controlled approach (92).

### Soft agar colony formation assay

Six-well plates were first coated with 1.5 ml of 0.5% agar dissolved in complete growing medium, and then allowed to harden for 1 hour at room temperature. After the first layer was set, the top layer containing 1×10^4^ cells in 1.5 ml of 0.3% agar dissolved in complete growing medium was added on top and allowed to harden for 30 minutes at room temperature. Plates were kept under standard cell culturing conditions for 2 to 3 weeks, with the regular addition of 300 μl of complete growth medium on top, to avoid drying of the plates. When treatment with NMDi was performed, the compound was added to both agar layers and to the growth medium added on top, at a final concentration of 0.3 μM. Colonies were stained for 1 hour at room temperature with crystal violet 0.01% (Sigma Aldrich) dissolved in 20% methanol and de-stained by performing several washes with water, before taking pictures with a gel documentation system. Automatic counting of the number of colonies was done using ImageJ software.

### Transwell migration assay

Transwell migration assays were performed using 12-well 8 μm PET transwell inserts (Sarstedt). 1×10^4^ cells were resuspended in 500 μl of FCS-free growing medium and dispensed into the upper chamber of the insert, in duplicates. The bottom well was then filled with either complete growing medium containing 10% FCS or FCS-free growing medium, which served as a control. When NMDi treatment was required, the inhibitor was added to the medium in both compartments, at a final concentration of 0.3 μM. Plates were incubated at 37 °C for 20 hours. Next day, non-migrating cells were removed from the upper chamber with a cotton swab and the migrated cells in the bottom side of the membrane were stained for 20 minutes at room temperature with crystal violet 1% in 20% methanol. After several washes with water, the membranes were cut and mounted on glass slides. Pictures of 5 random fields were taken, at 10x magnification, and the total number of migrated cells was counted.

### Growth curves

For measuring growth rates, 6×10^4^ cells/well were seeded in 12 wells of a 24-well plate in complete growth medium. Starting on the first day after seeding, the number of cells/ml present in three wells was measured every day, during 4 consecutive days. This protocol was performed in biological triplicates.

### Mouse Xenograft experiments

For testing the *in vivo* growing capacities of the cell lines, 2×10^6^cells resuspended in Matrigel (Corning) were injected subcutaneously in the flank of NSG mice. In total, 17 mice were injected per cell line. Tumor growth was followed over time by measuring its volume with a caliper, three times per week. When the tumors reached a volume of 1500 mm^3^, or when animals had to be sacrificed because of their health condition, half of the tumor tissue was harvested in RNAlater (Sigma Aldrich) and stored at −80 °C (for RNA extraction), and the other half was collected, fixed with formalin, and embedded in paraffin for histological examination. For the Kaplan-Meier plot, the time required for the tumors reaching 750 mm^3^ was taken.

### Histopathological analysis of tumors

Primary tumors tissue blocks were routinely processed and stained with hematoxylin and eosin (HE) for histological examination. The primary tumors were assessed morphologically for mitotic activity and for tumor necrosis. Quantitative measurements were done digitally using QuPath 0.2.3 software. For measuring the mitotic activity, 10 high power fields of 0.237 mm^2^ each were selected for counting mitosis. The assessment was done in the area (field of view, FOV) with estimated highest mitotic activity, selecting 10 consecutive fields. FOV with necrosis were excluded from the assessment. For the quantification of tumor necrosis, the areas of tissue with and without necrosis were measured and the percentage of necrosis was calculated.

### Gelatin zymograms

Twenty μl of conditioned media collected from nearly confluent plates were mixed with 20 μl of 2x Tris-Glycine SDS sample buffer (NOVEX LC2676), in the absence of reducing agents, and incubated at room temperature for 10 minutes. Next, the samples were loaded onto the Novex™ 10% Zymogram Plus (Gelatin) Protein Gels (ZY00100BOX, Invitrogen) and the gel was run at 125V using Tris-Glycine SDS running buffer. After the run was complete, the gel was incubated in Novex Zymogram Renaturing Buffer (LC2670, Invitrogen) for 30 minutes at room temperature, followed by a 30 minute incubation in Novex Zymogram Developing Buffer (LC2671, Invitrogen). The buffer was then exchanged with fresh developing buffer and the gel was developed overnight at 37 °C. Next day, the gel was stained with Imperial™ Protein Stain (Thermo Fisher Scientific) for 2 hours at room temperature and then destained by several washes with water. Finally, gels were imaged with a gel documentation system.

### Ingenuity Pathway Analysis

The list of differentially expressed genes between tumor samples, together with their fold changes, was uploaded to the Ingenuity Pathway Analysis (IPA) software and a core analysis was performed. Only differentially expressed genes with a fold change smaller than −2 and bigger than 2, and with an FDR smaller than 0.01 were considered for this analysis. The core analysis provided the functional characterization of our dataset, from which affected diseases or biological functions were inferred. The software was used to create “Regulators effects networks”, which are functional networks that connect differentially expressed genes (DEGs) with both downstream effects and upstream regulators. The IPA software was also used to compare the DEGs and the predicted downstream effects of the RNAseq data from the tumor samples and the one of the cells in culture.

### Graphs and statistical analysis

All graphs and statistical analysis were done with GraphPad Prism 9 software.

## Abbreviations

(DEG): Differentially expressed genes
(IPA): ingenuity pathway analysis
(KD): knock down
(KO): knock out
(NMD): nonsense-mediated mRNA decay
(NMDi): NMD inhibitor
(PTC): premature termination codon
(RT-qPCR): reverse transcription followed by real time PCR
(RNAseq): RNA sequencing
(WB): western blot.

## Acknowledgements and Funding

We thank Marieke van de Ven and her team from the MCCA Intervention Unit of the Netherlands Cancer Institute for performing the xenograft experiments, and Sophie Braga Lagache, Natasha Buchs, and Manfred Heller from the PMSCF of the University of Bern for their excellent mass spectrometry services. We also thank Nicole Kleinschmidt and Karin Schranz for their excellent technical support.

This work was supported by the National Center of Competence in Research (NCCR) on RNA & Disease funded by the Swiss National Science Foundation (SNSF; grants 51NF40-141735, 182880, and 205601), by SNSF grants 310030B-182831 and 310030-204161 to O.M., and by the canton of Bern (University intramural funding to O.M.). Funding for open access charge: SNSF.

## Notes

### Competing Interest Statement

The authors have declared no competing interest.

