## Supplemental Information for "Inhibition of nonsense-mediated mRNA decay reduces the tumorigenicity of human fibrosarcoma cells"

### Supporting Information for
### Inhibition of nonsense-mediated mRNA decay reduces the tumorigenicity of human fibrosarcoma cells.

Sofia Nasif, Martino Colombo, Anne-Christine Uldry, Markus S. Schröder, Simone de Brot, and Oliver Mühlemann*

**This PDF file includes:**

Figures S1 to S4
Table S1

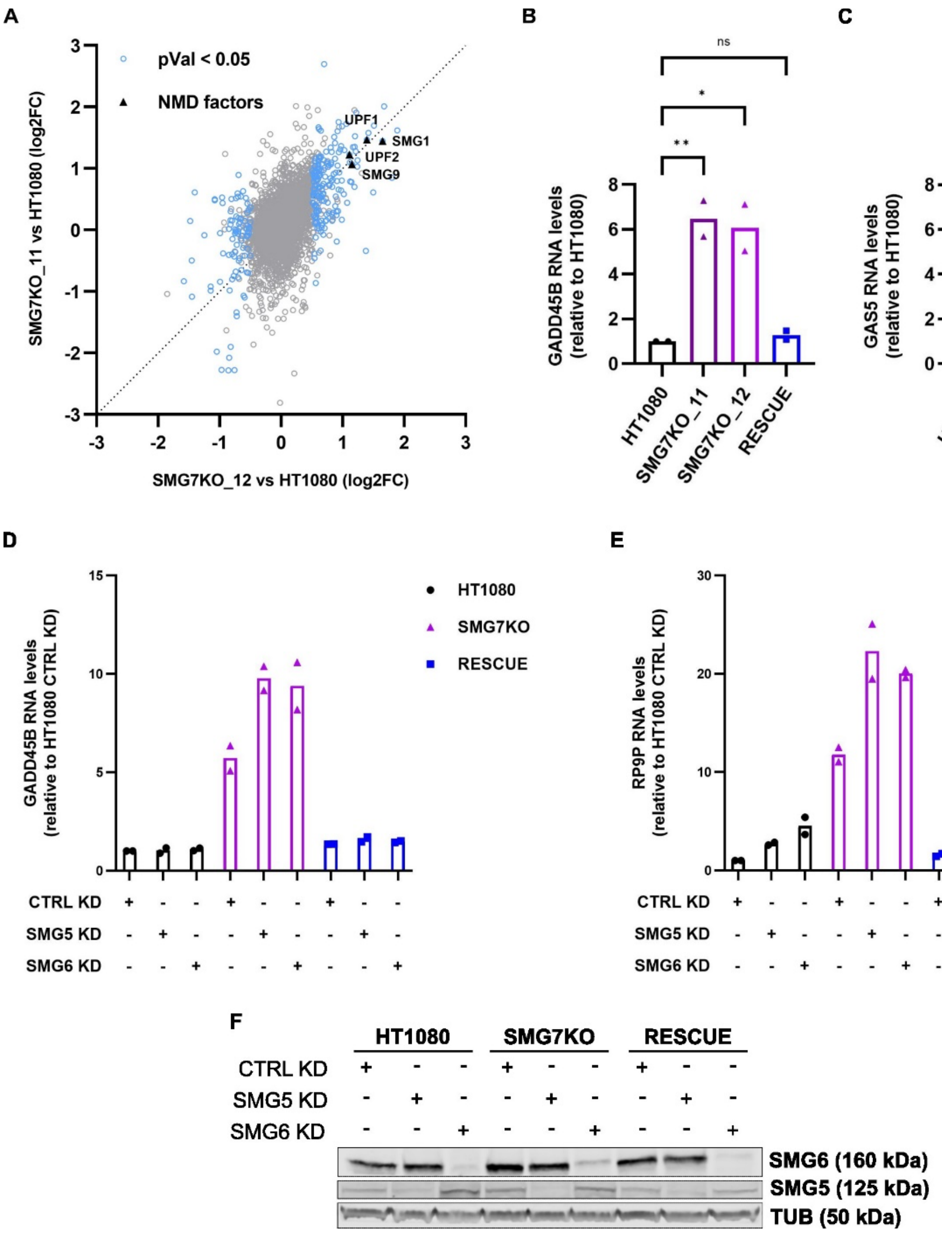


**Fig. S1.**

**(A)** Scatter plot showing the correlation between the differential protein expression analyses performed on SMG7KO clones 11 and 12 compared to the parental cell line HT1080. Differentially expressed proteins with *p*-values < 0.05 are shown in blue. The upregulated NMD factors are shown in black. **(B-C)** Relative mRNA levels, determined by RT-qPCR, normalized to ActB mRNA and 28S rRNA. Bars depict the mean values and data points represent results from biological replicates. Statistical significance was determined by One-way ANOVA and Dunnett's multiple comparisons test. *ns* p > 0.05; * p ≤ 0.05; ** p ≤ 0.01; **** p ≤ 0.0001. **(D-E)** Relative mRNA levels, determined by RT-qPCR, in HT1080, SMG7KO_12 and the RESCUE cell lines treated with the indicated siRNAs. mRNA levels were normalized to ActB mRNA and 28S rRNA levels. **(F)** Western blot showing the levels of SMG5 and SMG6 proteins in

HT1080, SMG7KO_12 and the rescue cell lines treated with the indicated siRNAs. Tubulin (TUB) served as a loading control.

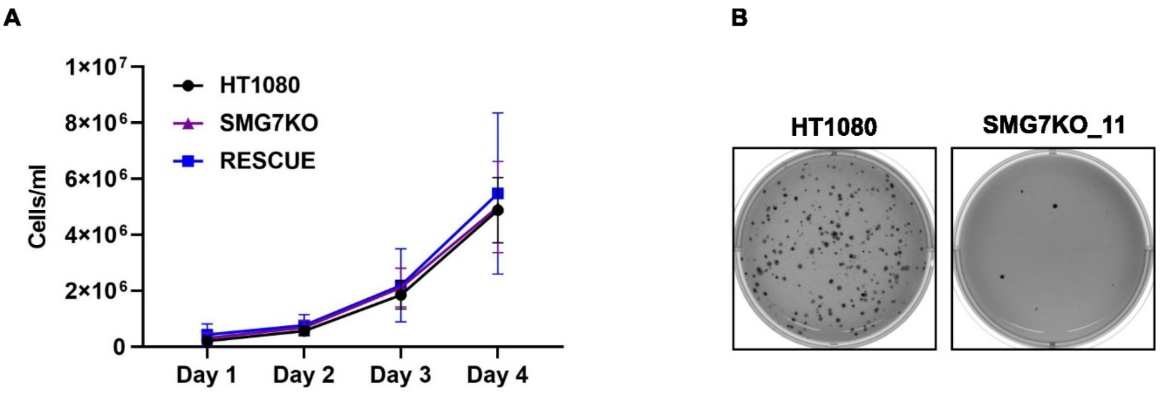


**Fig. S2.**

**(A)** Growth curves for HT1080, SMG7KO and RESCUE cell lines under standard cell culturing conditions. Shown are the means ± SD of three biological replicates. **(B)** Soft agar colony formation assays were used to determine the anchorage-independent growth properties of HT1080 and SMG7KO clone 11 cells.

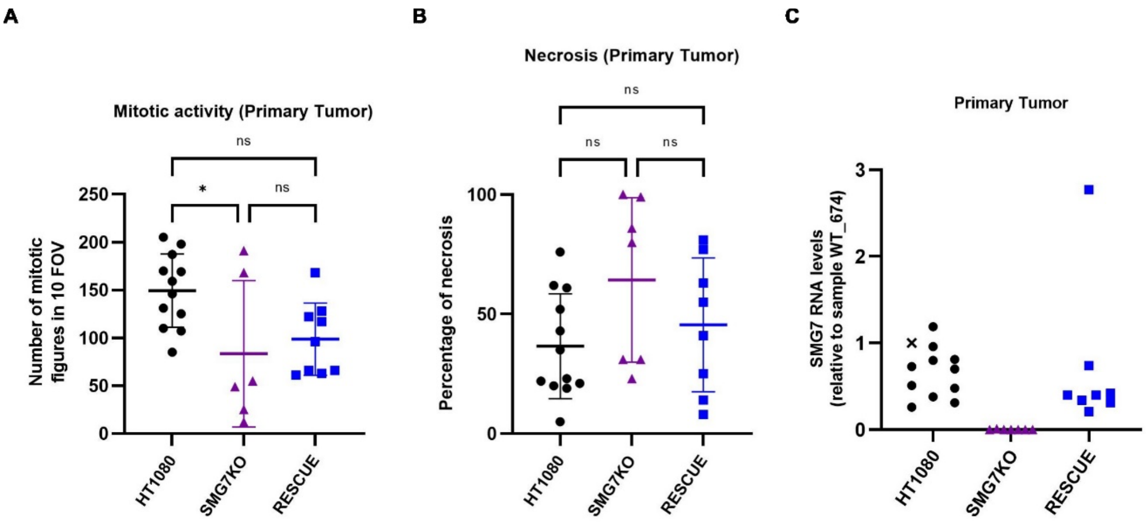


**Fig. S3.**

**(A)** Quantification of the mitotic activity in the primary tumors on hematoxylin and eosin stained tissue sections. Each data point represents the total number of mitoses found in 10 fields of view (FOV; $0.237mm^2$ per field) on each of the available primary tumor samples, HT1080 n = 12, SMG7KO n = 6 and RESCUE n = 9. Shown are the mean values ± SD, statistical significance was determined by One-way ANOVA and Tukey's multiple comparisons test. *ns* $p > 0.05$; * $p \leq 0.05$. **(B)** Quantification of necrosis in the primary tumors. Each data point represents the percentage of necrotic tissue (relative to the total area of tissue) on each of the available primary tumor samples, HT1080 n = 12, SMG7KO n = 7 and RESCUE n = 8. Shown are the mean values ± SD, statistical significance was determined by One-way ANOVA and Tukey's multiple comparisons test. *ns* $p > 0.05$. **(C)** Relative SMG7 mRNA levels, determined by RT-qPCR, normalized to ActB mRNA. Each data point represents a different primary tumor, and the values are expressed relative to the tumor sample HT1080_674, marked with a "x".

| Diseases or Functions Annotation | p-value | z-score |
|---|---|---|
| Invasion of cells | 8,64E-04 | -3,054 |
| Advanced malignant tumor | 1,24E-04 | -2,624 |
| Metastasis | 8,26E-04 | -2,624 |
| Invasive tumor | 1,48E-03 | -2,624 |
| Quantity of metal | 2,53E-03 | -2,196 |
| Incidence of tumor | 4,03E-06 | -2,166 |
| Migration of cells | 5,49E-05 | -2,119 |
| Arrhythmia of heart ventricle | 2,43E-03 | 2,000 |
| Inflammation of absolute anatomical region | 8,45E-04 | 2,107 |

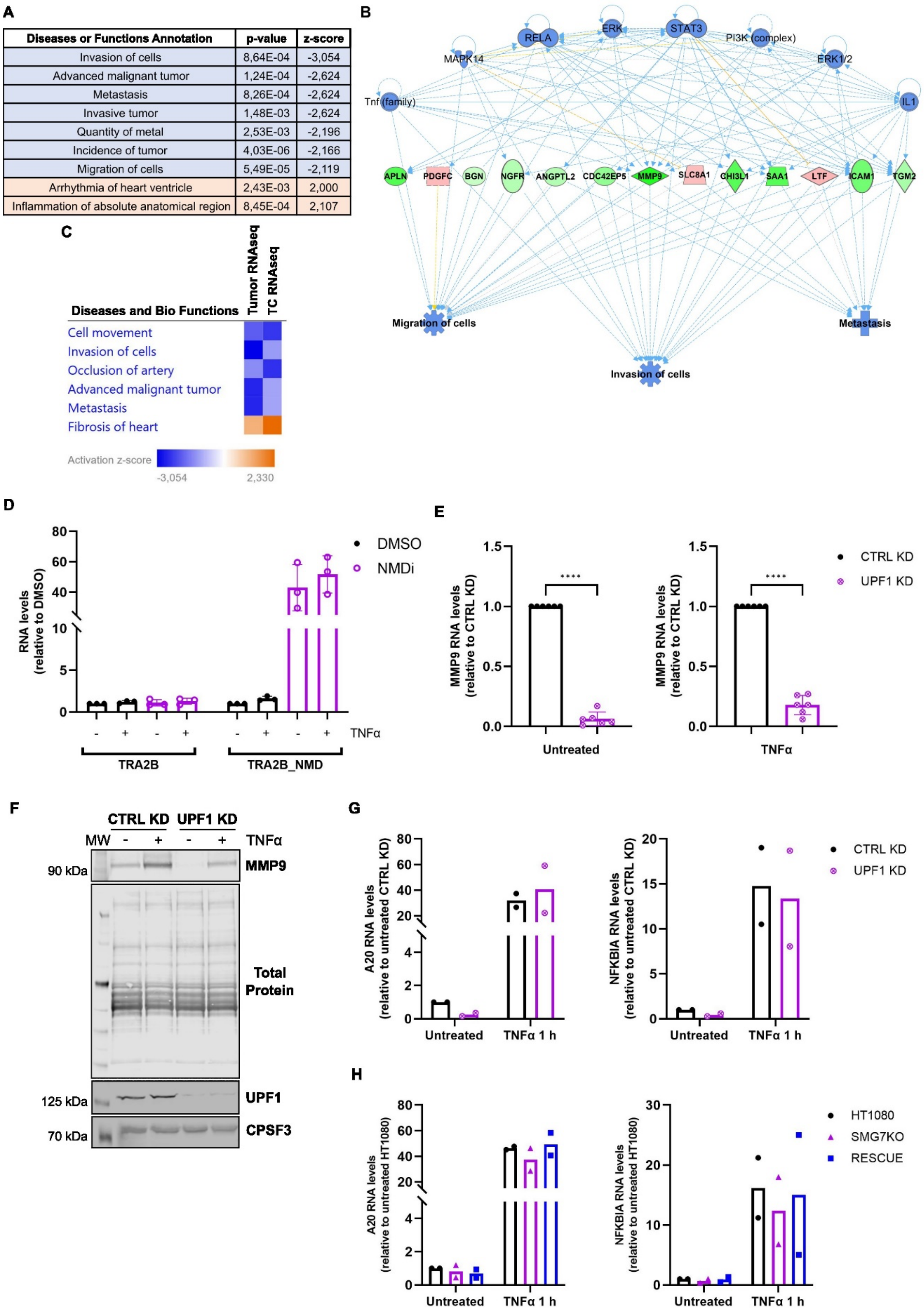

**Fig. S4.**

**(A)** The downstream effects analysis tool from IPA was used to analyze the RNAseq data from the primary tumors. The table depicts the diseases or biological functions predicted to be inhibited (z-score <-2, shown in blue) or activated (z-score >2, shown in orange) in the tumors formed by SMG7KO cells, compared to the tumors from HT1080 cells. **(B)** Regulators effects network created with the IPA software, using the RNAseq data from the primary tumors as input. Upstream regulators are shown in the top tier, their blue color shows they are predicted to be inhibited in this dataset. The bottom tier shows in blue the diseases or biological functions predicted to be inhibited, based on the gene expression changes observed in the dataset. The middle tier contains the genes that are suggested to mediate the effects between the upstream regulators and the downstream outcomes; downregulated genes are shown in green, upregulated ones are shown in red (SMG7KO tumors vs HT1080 tumors). Direct relationships are shown with filled lines and indirect relationships are shown with dotted lines; relationships leading to activation of the downstream node are shown in orange, if leading to inhibition they are shown in blue, and if the state of the downstream node doesn't match the prediction, they are shown in yellow. This causal network has a consistency score of 31.537. **(C)** Comparison analysis done with IPA on the RNAseq data from the primary tumors (Tumor RNAseq) and that of the cells in culture (TC RNAseq). The heat map shows the downstream diseases or functions predicted to be inhibited (negative z-scores, in blue) or activated (positive z-scores, in orange) in both datasets. Only the top six categories (based on their activation z-score) are shown. **(D)** Relative mRNA levels in HT1080 cells treated with 0.3 µM NMD inhibitor (NMDi) or with DMSO in the absence or presence of 25 ng/ml TNFα. mRNA expression was determined by RT-qPCR, normalized to ActB mRNA and 28S rRNA. Shown are the mean values ± SD and data points represent biological replicates. **(E)** Relative MMP9 mRNA levels in HT1080 cells treated with a control siRNA (CTRL KD) or an siRNA against UPF1 (UPF1 KD), in the absence or presence of 25 ng/ml TNFα. MMP9 expression was determined by RT-qPCR as (D). Statistical significance was determined by unpaired two-tailed t-test. **** $p \leq 0.0001$. **(F)** Western blot showing the levels of MMP9 protein present in conditioned medium collected from HT1080 cells undergoing a CTRL KD or a UPF1 KD, in the absence or presence of 25 ng/ml TNFα. Total protein staining serves as a loading control. Lower panel: Western blot was used to analyze UPF1 protein levels, CPSF3 serves as a loading control. MW: molecular weight marker. **(G)** Relative mRNA levels in HT1080 cells treated with a control siRNA (CTRL KD) or an siRNA against UPF1 (UPF1 KD), either untreated or after 1 hour stimulation with 25 ng/ml TNFα. Relative mRNA levels were determined as in (D). **(H)** Relative mRNA levels in HT1080, SMG7KO and RESCUE cells, untreated or after 1 hour stimulation with 25 ng/ml TNFα. Relative mRNA levels were determined as in (D).

**Table S1.** Antibodies and oligos

| Primary antibodies | | | | |
|---|---|---|---|---|
| **Target protein** | **Company** | **Cat N** | **Species** | **Dilution** |
| SMG7 | My BioSource | MBS820862 | Rabbit | 1/1000 |
| CPSF3 | Santa Cruz Biotechnology | sc-393001 | Mouse | 1/500 |
| Tyrosin-tubulin | Sigma Aldrich | T9028 | Mouse | 1/10000 |
| MMP9 | R&D systems | MAB911 | Mouse | 1/500 |
| FLAG | Antibodies online | ABIN99294 | Rabbit | 1/1000 |
| SMG5 | Abcam | ab33033 | Rabbit | 1/1000 |
| SMG6 | Abcam | ab87539 | Rabbit | 1/1000 |
| UPF1 | NeoBiotechnologies | 5976-MSM1-PAX | Mouse | 1/1000 |
| **Secondary antibodies** | | | | |
| **Target protein** | **Company** | **Cat N** | **Species** | **Dilution** |
| 2°-anti Mouse (green) | LI-COR Biosciences | IRDye® 800CW, 926-32212 | Donkey | 1/10000 |
| 2°-anti Mouse (red) | LI-COR Biosciences | IRDye® 680LT, 926-68022 | Donkey | 1/10000 |
| 2°-anti Rabbit (green) | LI-COR Biosciences | IRDye® 800CW, 926-32213 | Donkey | 1/10000 |
| 2°-anti Rabbit (red) | LI-COR Biosciences | IRDye® 680LT, 926-68023 | Donkey | 1/10000 |

| qPCR Oligos | | | |
|---|---|---|---|
| **Target** | **Forward primer** | **Reverse primer** | **Probe** |
| 28S rRNA | AGAGGTAAACGGGTGGGGTC | GGGGTCGGGAGGAACGG | |
| A20 | GCCATTGTAGTTGGTAGCCTTCA | TGCGCTGGCTCGATCTCAGTTG | |
| ACTIN | TCCATCATGAAGTGTGACGT | TACTCCTGCTTGCTGATCCAC | |
| GADD45B | TCAACATCGTGCGGGTGTCG | CCCGGCTTTCTTCGCAGTAG | |
| GAS5 | GCACCTTATGGACAGTTG | GGAGCAGAACCATTAAGC | |
| MMP9 | GAACCAATCTCACCGACAGG | GCCACCCGAGTGTAACCATA | |
| NFKBIA | CTCCGAGACTTTCGAGGAAATAC | GCCATTGTAGTTGGTAGCCTTCA | |
| RP9P | CAAGCGCCTGGAGTCCTTAA | AGGAGGTTTTTCATAACTCGTGATCT | |
| SMG7 | CAAGGCCAGGCAAAGAATCG | GGGAAGACTTCACACGGCAT | |
| SMG7 (TaqMan) | TCTTCCGTCCAGAGCAGGAT | AGCTCTGAGGGCTTCTCCAAT | FAM-CTGTACCCAGAATGCCGTTTGAGAAATCC-BHQ1 |
| TRA2B (protein coding, NMD insensitive) | GAGGTTGGCAGCTTCGATT | AAGCAGAACGGGATTCCC | |
| TRA2B_NMD (NMD sensitive) | TGGAATCAGAAAGCACTACGC | GAATCTTCCTTGGAGCGAGA | |

| siRNA Oligos | |
|---|---|
| **Target** | **Sequence** |
| Control | AGGUAGUGUAAUCGCCUUGdTdT |
| SMG5 | GAAGGAAAUUGGUUGAUACdTdT |
| SMG6 | GCTGCAGGTTACTTACAAGdTdT |
| UPF1 | GAUGCAGUUCCGCUCCAUUdTdT |
